# Feminization of sex-biased gene expression in a parthenogenetic stick insect suggests unresolved sexual antagonism in early development

**DOI:** 10.1101/2025.11.30.691440

**Authors:** Jelisaveta Djordjevic, Marjorie Labédan, Tanja Schwander, Darren J. Parker

## Abstract

Sexual antagonism arises when males and females experience opposing selection on traits controlled by shared genetic architectures. For gene expression, such conflicts are typically resolved through the evolution of sex-biased regulation. Although sexual antagonism is often assumed to peak in adults, its developmental dynamics remain poorly understood. We investigated the ontogeny of sex-biased gene expression in the sexual stick insect *Timema poppense* and contrasted it with its parthenogenetic sister species to identify signatures of unresolved antagonism. Using RNA-seq across somatic tissues (brain, gut, legs, antennae) and reproductive tracts throughout post-embryonic development, we found that sex-biased expression in *T. poppense* is limited in somatic tissues but extensive in reproductive tracts from the earliest stages. Nearly half of the expressed genes are already sex-biased in first-instar gonads, with male- and female-biased genes largely stage-specific. Parthenogenetic females show masculinized expression in somatic tissues throughout development and in adult reproductive tracts, consistent with decay of sexual traits in the absence of mating. By contrast, they show feminized early gonadal expression, likely reflecting unresolved antagonism in the sexual ancestor. Our results reveal the ontogeny of sexual dimorphism in a hemimetabolous insect and suggest that focusing solely on adults underestimates the prevalence of unresolved sexual antagonism.

## Introduction

Males and females have often diverged fitness optima for traits, leading to opposing selective pressures on these traits in the two sexes [1]. When, as a consequence of a largely shared genome, these traits have the same genetic architectures in males and females, sexual antagonism (also referred to as sexual conflict) emerges [2,3]. In the case of gene expression levels, antagonism over optimal expression levels can drive the evolution of different expression levels in males and females via sex-specific regulatory architectures. Sex-biased genes would thus reflect resolved or partially resolved sexual conflict over expression levels [4,5].

Although sexual conflict is predicted to be the strongest at the adult stage when sexual dimorphism is completely manifested [6,7], little is known about how sexual conflict and sex-biased gene expression vary over ontogeny [8]. Studies that investigated expression over ontogeny suggest that sex-biased gene expression is dynamic, and the extent of the bias depends on the analyzed tissue [9]. Thus, in *Timema* stick insects and *Nasonia* wasps, analyses of whole bodies show a general increase in sex-bias during development, mirroring the gradual increase of morphological sexual dimorphism [10,11]. Reproductive tracts in *Drosophila melanogaster* flies have a consistent sex-bias of approximately half of the genes from 3^rd^ instar larvae to the adult stage [8], while mammalian somatic organs show little sex-bias during development with an increase around sexual maturity [12]. These findings suggest that resolved but perhaps also unresolved sexual conflict over gene expression is present during development, and that its extent varies depending on the levels of sexual dimorphism of tissue functions.

The expression level of a typical gene is likely influenced by many loci and subject to stabilizing selection [13–15]. Recent theoretical approaches have shown that genetic polymorphisms affecting such polygenic traits under stabilizing selection are quickly lost, which makes the detection of unresolved sexual conflict at the level of individual genes difficult [16]. However, large-scale shifts in gene expression across the transcriptome can reveal the cumulative signature of such conflicts when sex-specific selection changes [17,18]. For instance, in obligately parthenogenetic species, where the male phenotype is no longer under selection, putative unresolved sexual conflict over gene expression disappears [19]. Consequently, genes that experienced opposing selection pressures in sexual ancestors are expected to shift their expression toward the female optimum, resulting in a feminized transcriptome in parthenogenetic females. This expectation has been probed in three different taxa, in *Timema* stick insects, *Artemia* brine shrimp, and *Acyrthosiphon* aphids [19–21]. However, contrary to expectations, parthenogenetic females were masculinized in stick insects and brine shrimps, i.e. had reduced expression of female-biased and greater expression of male-biased genes [19,20]. In the stick insects, convergent masculinization across independent parthenogenetic species was suggested to stem from vestigialized female sexual traits [22], which would mask potentially weaker signals of released sexual conflict [19]. In brine shrimp, masculinization was observed in sexual and parthenogenetic species relative to a shared outgroup, suggesting lineage-specific transcriptome shifts independently of reproductive mode [20]. In aphids, feminization of the transcriptome was observed for rare males from obligately parthenogenetic lines, but not females, perhaps because cyclical parthenogenesis in sexual lines already generates strong selection for adaptations to parthenogenesis [21]. However, because all three studies focused on adults and partially on composite tissues, it remains possible that unresolved conflict is masked by averaging gene expression across tissues or is more pervasive earlier in development.

Here we examine the developmental gene expression patterns in two closely related stick insect species, sexually reproducing *Timema poppense* and parthenogenetic *T. douglasi*. In these stick insects, there are no sex-specific genome regions (XX:XO sex determination; [23]), meaning that differences between the sexes stem solely from differential gene expression and post-transcriptional regulation. Note that genomic signatures in some all-female populations of *T. douglasi* indicate rare sexual reproduction [24]. However, rare sex in mostly parthenogenetic species does not change the prediction of selection on males being strongly relaxed as compared to females. Using the two *Timema* species, we investigate in depth the dynamics of sex-biased gene expression in somatic tissues, including the brain, legs, guts, and antennas, as well as in the reproductive tracts. We sequence RNA from those tissues across multiple post-embryonic developmental stages starting from the 1^st^ nymphal until the adult stage, whereby the ontogeny of *Timema* males and females consists of six and seven distinct stages, respectively [11]. Firstly, we explore sex-biased gene expression within the sexual species, the consistency of sex-bias across development, autosomal *vs* X-linkage of sex-biased genes, and gene networks associated with different tissues. This is followed by an examination of how sex-biased genes differ in expression between sexual and parthenogenetic females and whether we can identify signatures of unresolved sexual conflict. Finally, we also provide a detailed gene expression comparison between sexual and parthenogenetic females, across tissues and development.

## Material and Methods

### Insect husbandry

First instars were derived from eggs laid by captive-bred individuals initially collected in California in 2017. Our objective was to acquire 4 biological replicates from five distinct tissues (reproductive tracts and four somatic tissues) for each of seven developmental stages in sexual and parthenogenetic females (nymphal stages 1-6 and adults) and six stages in sexual males (nymphal stages 1-5 and adults; see Supplemental Table S1).

Upon hatching, insects were reared in petri dishes containing *Ceanothus* plant cuttings enveloped in moist cotton. They were nurtured until they reached a specific developmental stage, at which point they were dissected within 12h after molting. To track molting, we applied red acrylic paint to the thoraxes after each molt and checked daily for individuals lacking the paint. Prior to dissection, the insects were anesthetized using CO_2_.

From each developmental stage, we dissected the brain, antennae, legs, and guts. We initially only dissected reproductive tracts from the 4th stage onwards, as we encountered difficulties in unequivocally identifying reproductive tissues in earlier stages. After enhancing our dissection techniques, we were able to identify and dissect gonad tissues at earlier stages, and consequently, we included reproductive tract samples from newly hatched individuals (1st nymphal stage) at a later date.

During the dissection process, tissues were carefully placed into Eppendorf tubes containing ceramic beads and rapidly flash-frozen before being stored at -80°C for approximately one-third of the dissections. For the remaining dissections, due to laboratory closures during the pandemic, tissues were preserved in RNA later (Qiagen) before being stored at -80°C. Finally, for adult tissues, we used data from females from the same populations but which were dissected and sequenced for another project [25] (see Supplemental Table S1 for sample details).

### RNA extraction and sequencing

To each tissue-containing tube, 1 mL of TRIzol solution was added. Tissue homogenization was carried out using the Precellys Evolution tissue homogenizer (Bertin Technologies). Subsequently, 200 μL of chloroform was added to each sample, followed by vortexing for 15 seconds. The samples were then subjected to centrifugation at 12,000 revolutions per minute (rpm) at 4°C for 25 minutes. The upper phase, which contains the RNA, was transferred to a fresh 1.5 mL tube along with the addition of 650 μL isopropanol and 1 μL of Glycogen blue (GlycoBlue™ Coprecipitant). The samples were vortexed and placed at -20°C overnight. Samples were then centrifuged for 30 minutes at 12,000 rpm at 4°C. The liquid supernatants were removed, and the RNA pellet underwent two washes using 80% and 70% ethanol. Each wash was followed by a 5-minute centrifugation step at 12,000 rpm. Finally, the RNA pellet was resuspended in nuclease-free water and quantified using a fluorescent RNA-binding dye (QuantiFluor RNA System) and a nanodrop (DS-11 FX).

Library preparation using NEBNext (New England BioLabs) and sequencing on an Illumina NovaSeq 6000 platform with 100 bp paired-end sequencing (∼30 million reads per sample) was outsourced to a sequencing facility (Fasteris, Geneva).

### Quality control, mapping and counting

RNAseq reads were quality trimmed with trimmomatic [26] (v. 0.39, options: ILLUMINACLIP:AllIllumina-PEadapters.fa:3:25:6 LEADING:9 TRAILING:9 SLIDINGWINDOW:4:15). Any reads that were less than 80 bp long following trimming were discarded along with the corresponding read pair. Reads were then mapped to the *T. poppensis* genome PRJNA1126215 from [27] with STAR [28] (v 2.7.8a) with default settings except for the addition of two-pass mapping (--twopassMode Basic). Read counts were then obtained using HTseq [29] (v.011.2, options: --order=pos --type=exon --idattr=gene_id --stranded=reverse).

### Differential gene expression analyses

We assessed differential gene expression between the sexes using edgeR v.3.42.4 [30,31] in R Statistical Software v. 4.3.1 [32]. The analyses employed a negative binomial distribution-based generalized linear model and TMM normalization to determine p-values for gene count similarities among experimental groups. Data were analyzed separately for each tissue at each developmental stage (and only if at least two replicates per sex were available for given stage and tissue following QC, see Supplemental Table S1.). We filtered out genes with low counts, requiring a gene to be expressed in the majority of male or female libraries, dependent on the number of replicates per sex (i.e., at least three libraries for four replicates, and two libraries for three replicates), with an expression level above 0.5 CPM.

Differential gene expression between sexes was examined using a generalized linear model with a quasi-likelihood F-test [33]. Multiple testing was corrected using the Benjamini-Hochberg method [34] with a 5% significance level. Genes with significantly higher expression in males were considered male-biased, genes with higher expression in females were labeled female-biased, and genes with no significant differential expression were categorized as unbiased. Genes were further classified into categories depending on the strength of bias into; slight (< 2 FC), strong (> 2 FC), or sex-limited (i.e., not expressed in opposite sex). To visualize the overlap of sex-biased genes between developmental stages we used ggVennDiagram by ggplot2 v.3.4.4 [35].

### Functional enrichment analysis

We performed Gene Ontology (GO) enrichment analysis using TopGO v.2.54.0 [36], on sex-biased genes identified with edgeR (as described earlier). Genes were functionally annotated by comparing the longest isoform of each gene to NCBI’s nr *Drosophila melanogaster* database (taxa id: 7227) using BlastP via Blast2GO [37] within OmicsBox [38] (v.3.1.2) with default options. Interproscan (OmicsBox default settings) was then run and merged with the BLAST results to obtain GO terms. Node size was set to ten, indicating the inclusion of only GO terms with annotations for at least 10 genes. Term enrichment was assessed using a weighted Kolmogorov-Smirnov-like statistical test, equivalent to the gene set enrichment analysis (GSEA) method, which considers all genes without applying an arbitrary threshold [39]. The “elim” algorithm was applied, considering the Gene Ontology hierarchy, prioritizing the assessment of the most specific GO terms before evaluating more general ones. Our analysis specifically focused on gene sets within the Biological Processes (BP) category of GO. Terms were deemed significant if the p-value was below 0.05.

### Expression differences between sexual and parthenogenetic females

We called differential gene expression between sexual and parthenogenetic females as described above for males and females of the sexual species. Next, to investigate whether genes with sex-biased expression in the sexual species (*T. poppense*) exhibited increased or decreased expression levels in parthenogenetic females, we compared expression levels for those genes between sexual and parthenogenetic females. We only used tissues and developmental stages with at least 20 sex-biased genes for these comparisons and used Wilcoxon rank sum tests to compare genes categorized as female-biased and male-biased, relative to genes with no sex-bias. To account for multiple testing, we applied False Discovery Rate (FDR) correction to the p-values obtained. We also repeated these analyses using a more robust set of genes by excluding genes with coefficients of variation (CV) in expression greater than one across all replicates. This filtering ensured that the comparisons focused on genes with more stable expression levels, reducing the influence of highly variable genes that might disproportionately contribute to apparent changes in expression.

## Results

### Sex-biased gene expression

Somatic and reproductive tissues differ in their dynamics of sex-biased gene expression during development. Somatic tissues have few sex-biased genes across all developmental stages, with an increase at the adult stage for a subset of tissues (**Figure 1**). The brain has the lowest percentage of sex-biased genes, ranging between 0 and 0.02 % across development, and without an increase at the adult stage (**Figure 1**). In guts, the percentage of sex-biased gene expression ranges between 0 and 0.4%, and in legs between 0 and 0.5%. Antennae have the greatest sex-bias, with 10% of genes being sex-biased at the adult stage. Sex-biased genes in each tissue are enriched for metabolic processes associated with physiological and molecular roles of that tissue (**Supplemental Table S2**).

The distribution of sex-biased genes differs between the X chromosome and the autosomes. In the antennae of adults, the X is feminized, i.e., significantly enriched for female-biased (Fisher’s exact test, padj < 0.01) and depleted for male-biased genes (Fisher’s exact test, padj < 0.01). By contrast, in legs, the X is enriched for both female and male-biased genes at the 3rd nymphal stage (Fisher’s exact test male-biased; padj.= 0.015, female-biased; padj.< 0.01). Sex-biased genes in the gut do not have a differential distribution between the X and autosomes, for neither male- or female-biased genes.

In the reproductive tract, the percentage of sex biased genes increases only slightly over development, with the extent of sex bias increasing more markedly. At the 1^st^ nymphal stage, 5651 (44%) of the expressed genes show sex-biased expression, with 28% of them displaying strong sex bias, characterized by a fold change (FC) > 2 (adjusted p-value <0.05) (**Figures 1, 2**). At the 4^th^ nymphal stage, 6619 (45%) of the genes exhibit sex-biased expression, with 39% of them displaying strong sex bias (FC > 2) (adjusted p-value <0.05) (**Figures 1, 2**). At the adult stage, 7748 (54%) of the genes are sex biased, with 46% of them exhibiting strong sex bias (FC > 2) (adjusted p-value <0.05).

The vast majority of male-biased and female-biased genes are stage specific (**Figure 3**). Only 278 (4%) male-biased genes remain male-biased in all three stages, while 748 (12%) remain female-biased (**Figure 3**). Furthermore, sex bias is not strongly skewed towards one sex, however the proportion of female- and male-biased genes differs significantly at every developmental stage. At the 1^st^ nymphal stage more genes are female biased with 24.2%, whereas 20% are male-biased (χ2(1) = 56.09, p = 6.92e-14). In later stages, there are more male-biased genes compared to female-biased ones. Specifically, at the 4th nymphal stage, 25.4% are male-biased and 19.4% are female-biased (χ2(1) = 118.87, p < 2.2e-16). In the adult stage, this trend continues with 30% of male-biased genes and 23.4% of female-biased genes (χ2(1) = 107.35, p < 2.2e-16).

The distribution of sex-bias on autosomes and the X chromosome in the reproductive tract is expected to differ from somatic tissues, given that *T. poppense* males silence their X chromosome during meiosis in germ cells [27]. Accordingly, the majority of X-linked genes are female-biased, 62% at 4^th^ nymphal and 66% at adult stage (**Figure 2 and Supplemental Figure S1**). At the first nymphal stage when the X is dosage compensated and not yet silenced [27], we detect 12% of genes on the X with male-bias and 29.5% with female bias. Female-biased genes are significantly enriched (Fisher’s exact test, p_adj._ = 0.007), and male-biased genes significantly depleted on the X at this stage (Fisher’s exact test, p_adj._ < 0.01).

**Figure 1.**
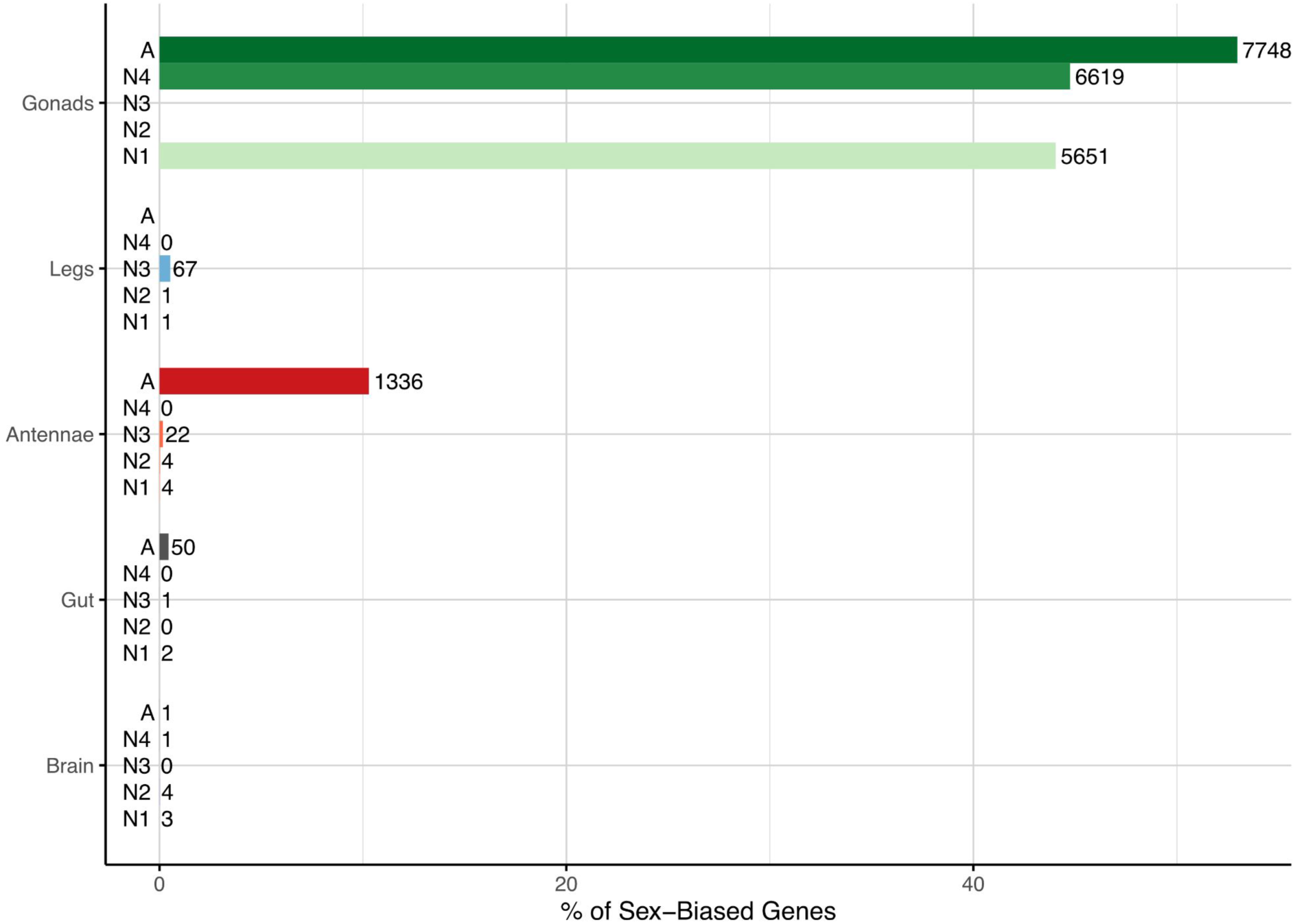
The percentage of sex-biased genes in somatic and reproductive tissues throughout development. The number of sex-biased genes is indicated next to each bar, with tissues color-coded (green-gonad, blue-legs, red-antennae, grey-gut, violet-brain) and shaded based on developmental stages from Nymphal stage 1 (N1) to Adult stage (A). The average number of expressed genes across tissues and developmental stages is 12,937 ± 824.35 (mean ± SD). Developmental stages are denoted as A (adult) and [N1-N4] (1st to 4th nymphal).

**Figure 2.**
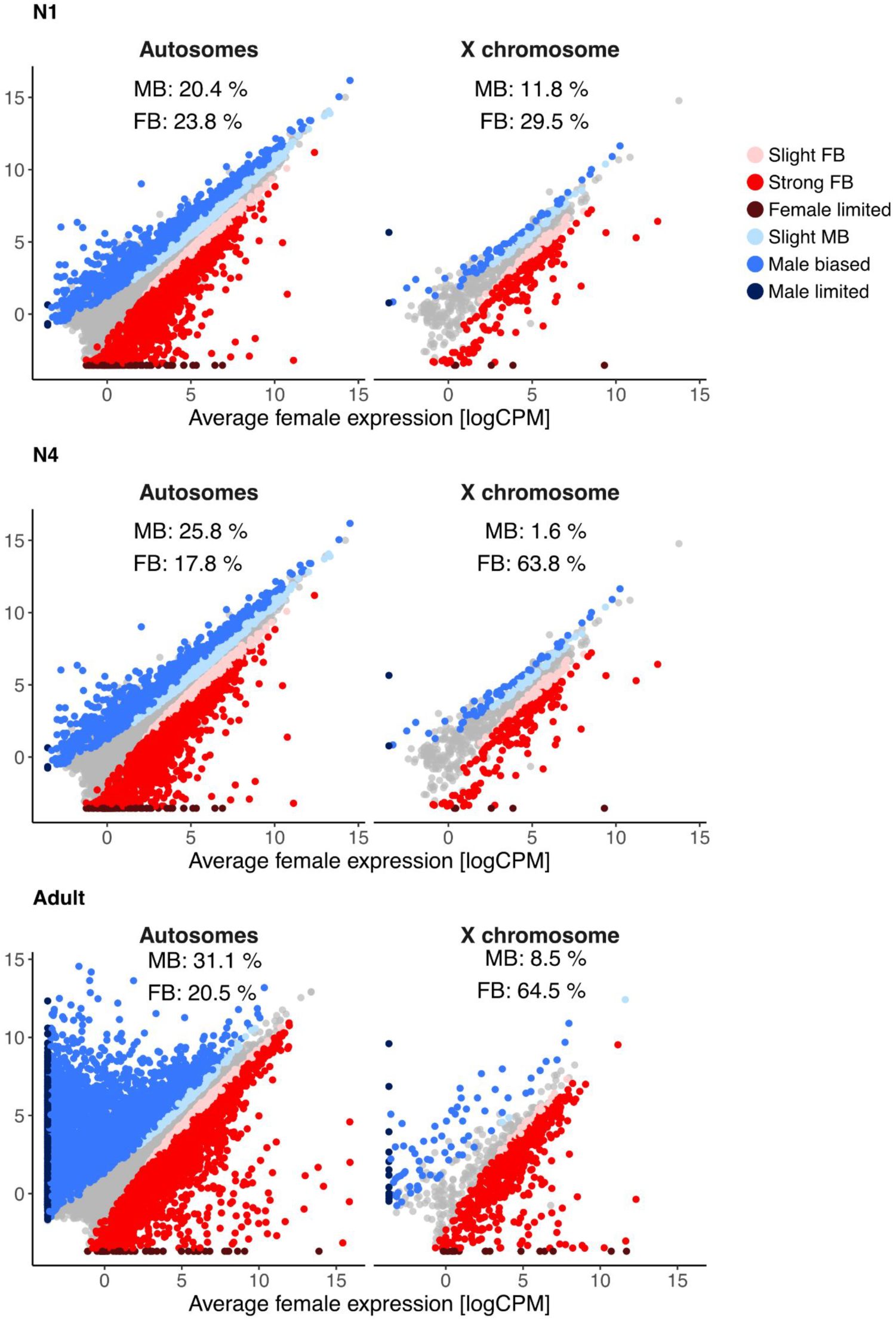
Sex-biased gene expression in the reproductive tract for three developmental stages. Panels on the left show expression on the autosomes, and right-side panels show expression on the X-chromosome. Genes are classified into categories depending on the strength of the sex-bias, slight female or male biased genes < 2 FC, strong female or male biased genes > 2 FC, female or male limited genes, have 0 expression the opposite sex, and un-biased genes are not significantly differentially expressed. Note that the presence of X-linked, male biased genes on the X is expected even though the X chromosome is silenced in male meiotic cells (which are present in all stages except N1 [27]). This is because male reproductive tracts comprise many somatic cells as well as germ cells that have not yet entered meiosis.

**Figure 3.**
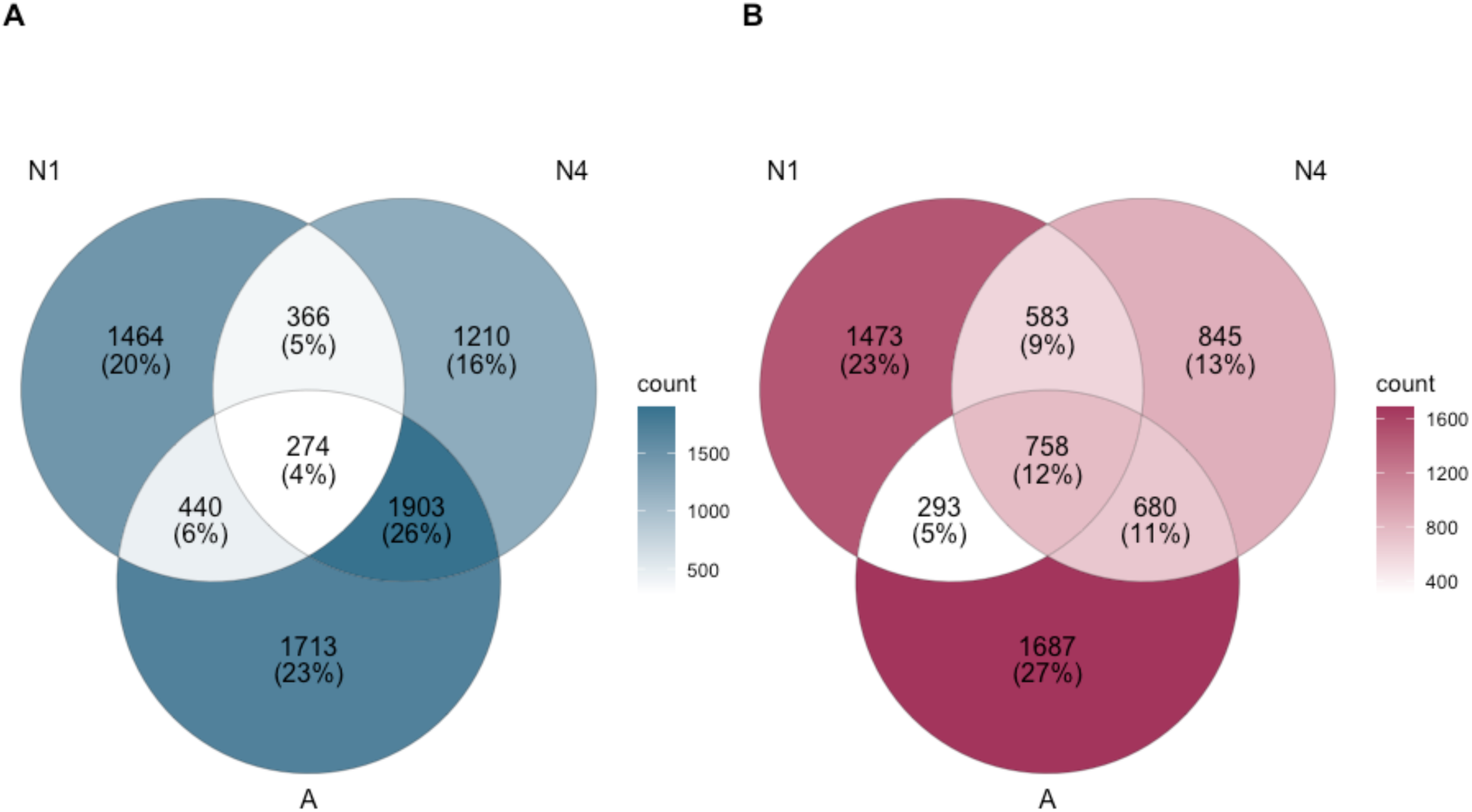
Venn diagrams showing the overlap of sex-biased genes from the reproductive tract between developmental stages **A)** Male-biased genes **B)** Female-biased genes

### Differential gene expression between sexual and parthenogenetic females

Sexual and parthenogenetic females generally feature strongly differentiated gene expression at the first nymphal and adult stages, and very similar expression throughout intermediate nymphal stages (**Figure 4**). This is the case for both somatic and reproductive tissues. Specifically, at the 1st nymphal stage, differential gene expression ranges between 13.4 and 39.6% for brain, gut, and reproductive tract, while legs and antennae have 0.8 and 0.4% of differentially expressed genes (DEG) respectively. Later nymphal stages feature very few DEGs, ranging between 0.1 and 1.2%. At the adult stage there is an increase with percentages of DEGs between 3.2 and 17.2%.

**Figure 4.**
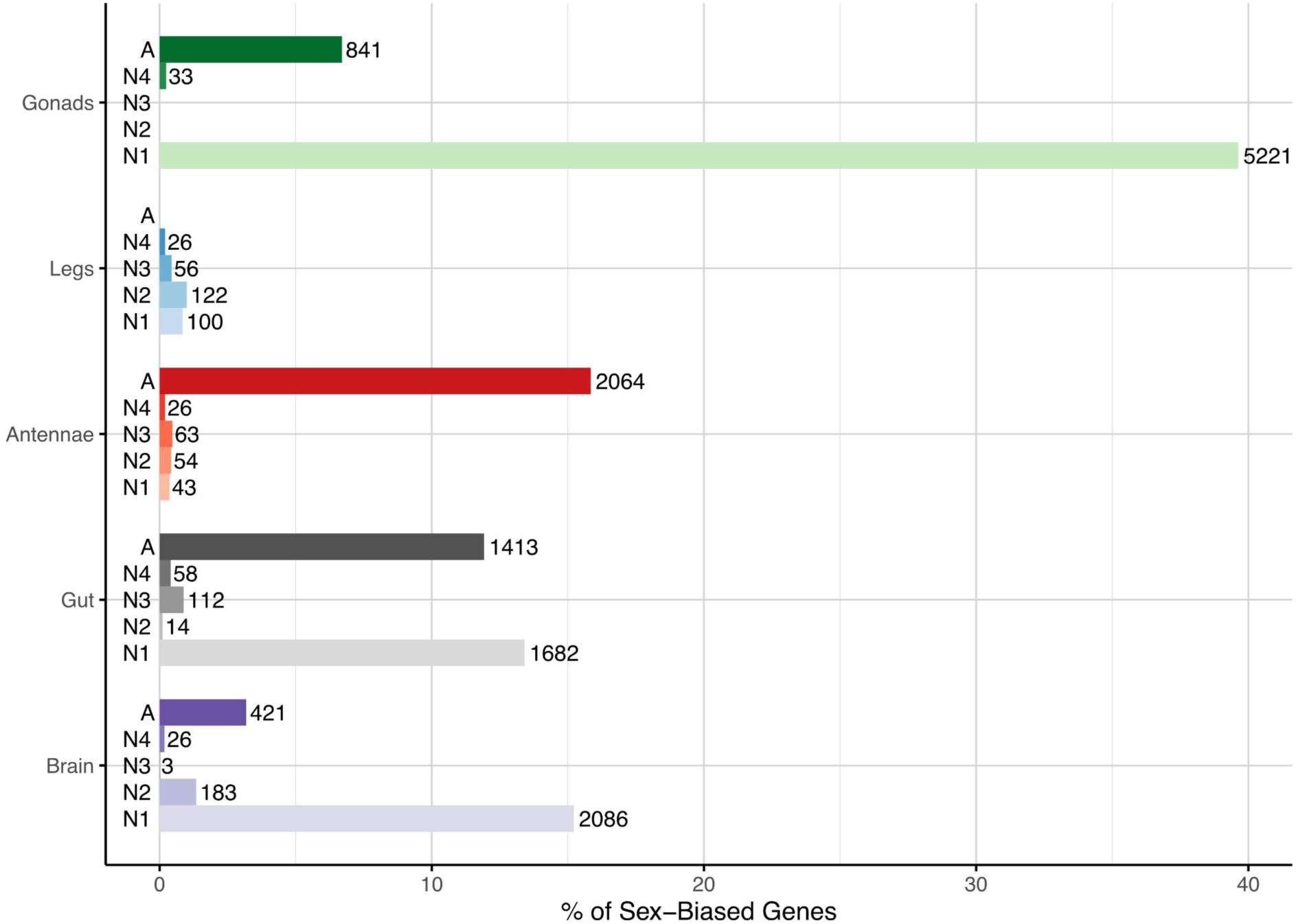
Percentage of the differentially expressed genes between sexual and parthenogenetic females in somatic and reproductive tissues, throughout development. The number of differentially expressed genes is indicated next to each bar, with tissues color-coded (green-gonad, blue-legs, red-antennae, grey-gut, violet-brain) and shaded based on developmental stages from Nymphal stage 1 (N1) to Adult stage (A). Average number of expressed genes across tissues and developmental stages: 12967 ± 711.7 (mean ± SD). Developmental stages are denoted as A (adult) and [N1-N4] (1st to 4th nymphal).

Finally, we examined differences between sexual and parthenogenetic females for genes with sex-biased expression in the sexual species (*T. poppense*). In the reproductive tract, 1^st^ nymphal stage parthenogenetic females are characterized by feminized gene expression: they have higher expression of female-biased genes, and lower expression of male-biased genes than sexual females (**Figure 5A**). We found the same pattern when analysing the subset of genes with little variation across biological replicates (**Supplemental Figure S2**). In the 4th nymphal and adult stages, we record opposite shifts in sex-biased gene expression, i.e. masculinization: higher expression of male-biased genes and lower expression of female biased genes in parthenogenetic females (**Figure 5A**). At all three stages, approximately half of the genes differentially expressed between sexual and parthenogenetic females are also sex-biased in the sexual species, independently of the considerable variation in differential expression between females at different developmental stages (33, 841, or 5221 genes, **Figure 5B**). Finally, similarly to the pattern observed in reproductive tracts at the 4^th^ nymphal and adult stages, parthenogenetic females show masculinization of expression in somatic tissues compared to sexually reproducing females (**Supplemental Figure S3**).

**Figure 5.**
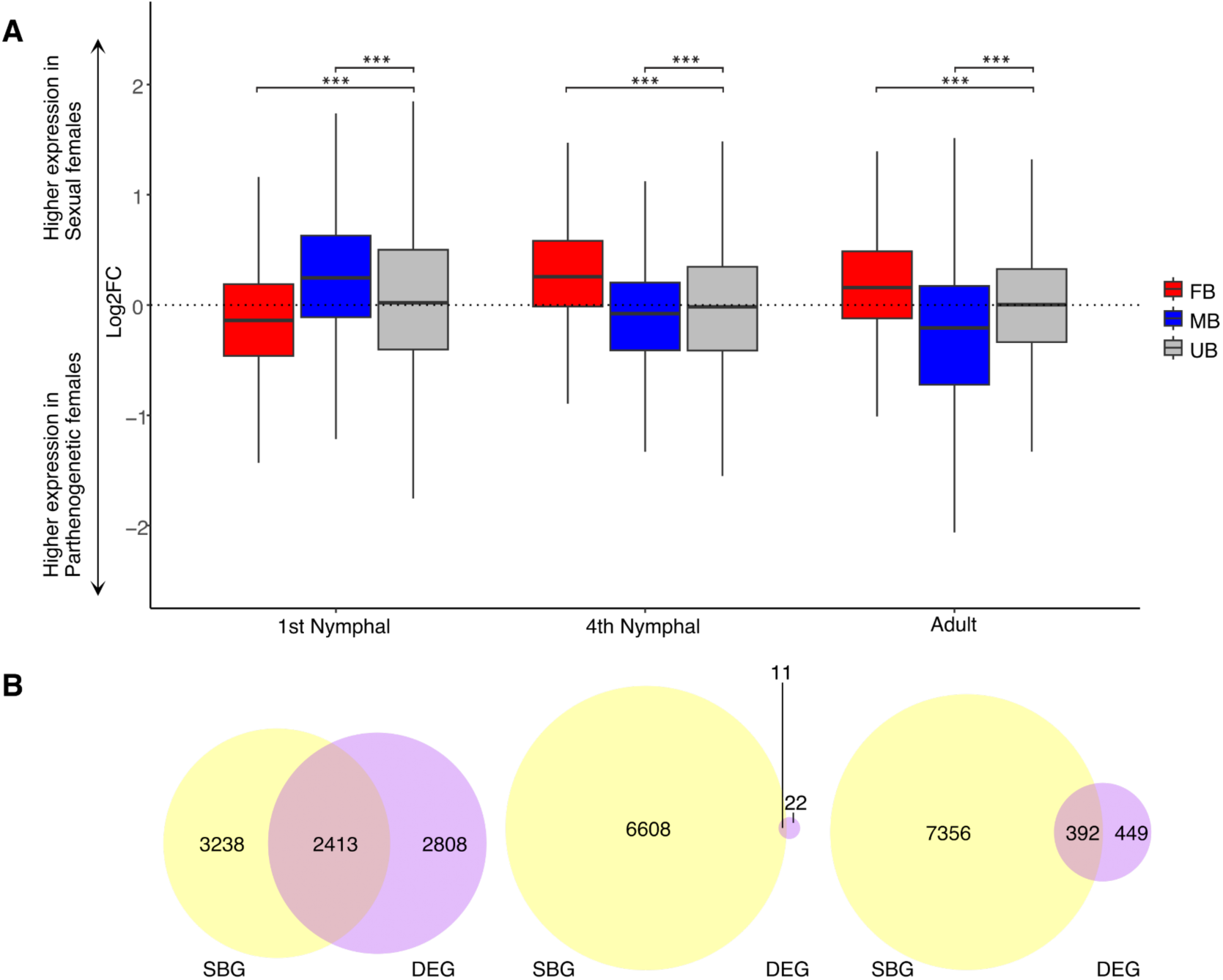
**A)** Shifts in expression levels of sex-biased genes between sexual and parthenogenetic females across development (in reproductive tracts). Three gene categories are depicted with different colors; female-biased in red, male-biased in blue, and un-biased in grey. Boxplots represent the median, lower and upper quartiles, and whiskers the minimum and maximum values (in the limit of 1.5x interquartile range). All comparisons between gene categories at each developmental stage yielded statistically significant results, as indicated by adjusted p values from Wilcoxon rank sum tests, with *** denoting p < 2e-16. **B)** Venn-diagrams showing overlaps between sex-biased genes (SBG) identified in the sexual species, and differentially expressed genes between sexual and parthenogenetic females (DEG) in reproductive tracts.

## Discussion

Our study reveals a stark contrast between somatic and reproductive tissues in the developmental dynamics of sex-biased gene expression in *Timema poppense*. While somatic tissues show little sex-bias throughout development, with the notable exception of adult antennae, the reproductive tract exhibits extensive and early sexual differentiation. Remarkably, already at the 1st nymphal stage, more than 40% of genes in the reproductive tract are sex-biased, a level that changes only slightly through to adulthood.

This trajectory differs from patterns described in *Drosophila melanogaster*, where approximately half of all genes are sex-biased from the third larval stage to adulthood, and a large proportion of these genes remain sex-biased throughout development [8]. In contrast, only a minority of sex-biased genes in *Timema* reproductive tracts are shared across developmental stages. The cause of this discrepancy remains unclear, as perhaps more similarity between tissues across stages in hemimetabolous (stick insects) than holometabolous (flies) insects could be expected [11]. It may reflect consequences of drift in stick insects, or species-specific timing of key reproductive processes, like ovary differentiation and egg provisioning, rather than fundamental differences between hemimetabolous and holometabolous development.

Our previous whole-body analyses in *T. californicum* suggested minimal sex-biased gene expression early in development [11]. Our current tissue-specific approach, however, reveals that this pattern is driven primarily by somatic tissues: reproductive tissue is already highly sexually differentiated at early stages, but its small proportional contribution to total body mass likely masked this effect in previous whole-body studies (see also [40]. This suggests that the observed gradual increase in sex-biased expression across development in *T. californicum* is largely attributable to the growth of reproductive tissues rather than a true shift in gene expression dynamics.

Among somatic tissues, the extent of sex-biased gene expression also varies. The brain shows almost no sex bias at any stage, with only 0.05% of expressed genes sex-biased even in adults. This pattern, also seen in fruit flies, crickets, and mammals [12,41,42], supports the idea that many sex-specific neural functions stem more from differences in circuitry than from transcriptional regulation. Notably, our whole-brain RNA-seq approach may also mask regional expression differences, as observed in other systems [40]. By contrast, adult antennae show substantial sex bias (10% of expressed genes), likely reflecting the development of sex-specific sensory roles, particularly in pheromone detection, a process expected only after sexual maturity [43,44]. The gut also displays modest sex-biased expression at the adult stage, enriched for genes involved in metabolic processes. These patterns suggest that even low levels of sexual dimorphism in somatic tissues, whether associated with diet [45], behavior [46,47], or niche use [48], can drive sex-specific gene regulation.

Patterns of sex-bias driven by X-linked vs autosomal genes can provide insights into the evolutionary forces shaping sex-specific expression [4,49,50]. As in many species (e.g., [8,51,52], female-biased genes in *T. poppense* are generally enriched on the X across tissues, while male-biased genes tend to be depleted. The overrepresentation of female-biased genes on the X chromosome is commonly attributed to the X being present twice as often in females as in males over the course of its evolution [53–55]. However, the observed depletion of male-biased genes on the X is less well understood. In somatic tissues, mechanistic limits imposed by dosage compensation, specifically how much a single X in males can be upregulated, have been proposed as a constraint [27,56]. In reproductive tissues, the inactivation of the X during male meiosis (MSCI) has been suggested as a major determinant of gene distribution (X or autosome) and sex-specific expression [57,58]. In *Timema* gonads, we here show that female-bias and depleted male-bias of the X persists throughout development, whereby the X is fully dosage-compensated in male gonads during early development (first nymphal stage) but becomes inactivated with the onset of meiosis [27]. In other words, the X is consistently feminized, independent of whether it is upregulated (via dosage compensation) or partially suppressed (via MSCI). This pattern strongly suggests that sexual antagonism, rather than mechanistic constraints on X upregulation or consequences of MSCI, is the key driver of gene distribution in this tissue.

Comparisons between sexual and parthenogenetic females further support the presence of unresolved sexual antagonism, particularly in early reproductive tissues. While adult parthenogenetic females show masculinized gene expression across tissues (as previously observed [19]), we detect the opposite pattern at the 1st nymphal stage in the reproductive tract: parthenogenetic females exhibit clear feminization, with increased expression of female-biased genes and decreased expression of male-biased genes. This suggests that sexual conflict over early gonadal gene expression is not fully resolved in the sexual species and is only revealed when male-specific selection stops in a parthenogenetic species. The absence of sexual trait decay in early-stage gonads likely facilitates the detection of this signal. Indeed, we do not expect to observe sexual trait decay in 1^st^ instar gonads in *Timema* as early oogenesis processes are likely very similar in parthenogenetic and sexual females. Both types of females produce an oocyte that contains a haploid maternal nucleus[59]. While in sexual females, diploidy of the embryo follows upon fertilization, diploidy of the parthenogenetic embryo occurs during development via gamete duplication[59,60]. Sexual conflict over gene expression in early gonads may be more difficult for selection to resolve at this early stage as the reproductive tracts start to extensively differentiate between the sexes. At the adult stage this tissue has very different morphology, function, and gene expression in each sex, to the point where testes and ovaries could almost be considered as two separate tissues (albeit with homologous origins). Testes and ovaries thus depend on very different regulation in each of the sexes meaning that any changes in gene expression in adults of one sex are unlikely to impact expression in the other.

## Conclusion

The dynamics of sex-biased gene expression differ extensively between somatic and reproductive tissues in *Timema poppense*. Somatic tissues have minimal sex-bias across development with an increase at the adult stage in only one of the tissues (antennae), while reproductive tracts show extensive differences from early development. Sexual and parthenogenetic females differ the most in gene expression at 1^st^ nymphal stage and in particular in the reproductive tract. At this stage, parthenogenetic females show feminization of gene expression in the reproductive tract, suggesting unresolved sexual conflict in early gonads of the sexual species. This supports the idea that sexual conflict can occur early in development, and is often overlooked by focusing only on adult stages.

## Acknowledgements

We thank Bart Zijlstra, Armand Yazdani, Susana Freitas, and Chloé Larose for help in the field, and current and previous members of the Schwander lab for discussions.

## Author contributions

J.D., T.S. and D.J.P., designed the study. J.D., and M.L. performed molecular work. J. D., analyzed the data with input from T.S.. and D.J.P.. J.D. wrote the paper with input from all authors

## Data and code availability

Raw sequence reads have been deposited in NCBI’s sequence read archive under the following bioproject: PRJNA1128554 (RNAseq reads). Data were processed to generate plots and statistics using R v3.4.4. Code is available here: https://github.com/JelisavetaDjordjevic/Sexual_conflict_development/ and will be archived upon acceptance.

