## Supplementary material for "Feminization of sex-biased gene expression in a parthenogenetic stick insect suggests unresolved sexual antagonism in early development": Supp material

### Supplemental Material

**Supplemental Table S1.** Final number of replicates for males (M), females (F) and asexual females (Asex F), per stage and tissue that passed quality control.

| | Adult | | | Nymphal 1 | | | Nymphal 2 | | | Nymphal 3 | | | Nymphal 4 | | | Nymphal 5 | | | Nymphal 6 | | |
|---|---|---|---|---|---|---|---|---|---|---|---|---|---|---|---|---|---|---|---|---|---|
| Tissue | Asex F | F | M | Asex F | F | M | Asex F | F | M | Asex F | F | M | Asex F | F | M | Asex F | F | M | Asex F | F | M |
| Antennae | 4 | 4 | 4 | 3 | 3 | 2 | 3 | 3 | 3 | 3 | 2 | 3 | 2 | 3 | 3 | 4 | 1 | 3 | 1 | 1 | 0 |
| Brain | 4 | 4 | 4 | 3 | 3 | 3 | 3 | 3 | 3 | 3 | 2 | 3 | 2 | 3 | 3 | 4 | 1 | 2 | 1 | 1 | 0 |
| Gonads | 4 | 4 | 3 | 4 | 4 | 4 | 0 | 0 | 0 | 3 | 2 | 1 | 2 | 3 | 3 | 3 | 1 | 3 | 1 | 1 | 0 |
| Guts | 4 | 4 | 4 | 3 | 3 | 3 | 3 | 3 | 3 | 3 | 2 | 3 | 2 | 2 | 3 | 2 | 0 | 2 | 1 | 1 | 0 |
| Legs | 0 | 0 | 0 | 3 | 3 | 3 | 3 | 3 | 3 | 3 | 2 | 3 | 2 | 3 | 3 | 4 | 1 | 3 | 1 | 1 | 0 |

**Supplemental Table S2.** Go term enrichment of sex-biased genes for different tissues and developmental stages

| Gonad N1 | |
|---|---|
| GO:0000398 | mRNA splicing, via spliceosome |
| GO:0048284 | organelle fusion |
| GO:0051173 | positive regulation of nitrogen compound metabolic process |
| GO:0006364 | rRNA processing |
| GO:0044260 | cellular macromolecule metabolic process |
| GO:0031325 | positive regulation of cellular metabolic process |
| GO:0051246 | regulation of protein metabolic process |
| GO:0007281 | germ cell development |
| GO:0051240 | positive regulation of multicellular organismal process |
| GO:0051172 | negative regulation of nitrogen compound metabolic process |

| Gonad N4 | |
|---|---|
| GO:0010996 | response to auditory stimulus |
| GO:0003341 | cilium movement |
| GO:0060271 | cilium assembly |
| GO:0072528 | pyrimidine-containing compound biosynthetic process |
| GO:0071407 | cellular response to organic cyclic compound |
| GO:0045892 | negative regulation of DNA-templated transcription |
| GO:0010970 | transport along microtubule |
| GO:0097305 | response to alcohol |
| GO:0035883 | enteroendocrine cell differentiation |
| GO:0000278 | mitotic cell cycle |

| Gonad A | |
|---|---|
| GO:0003341 | cilium movement |
| GO:0035336 | long-chain fatty-acyl-CoA metabolic process |
| GO:0048663 | neuron fate commitment |
| GO:0043266 | regulation of potassium ion transport |
| GO:0035046 | pronuclear migration |
| GO:0071577 | zinc ion transmembrane transport |
| GO:0010996 | response to auditory stimulus |
| GO:0042335 | cuticle development |
| GO:0048512 | circadian behavior |
| GO:0043171 | peptide catabolic process |

| Antenna N3 | |
|---|---|
| GO:0045752 | positive regulation of Toll signaling pathway |
| GO:0040003 | chitin-based cuticle development |
| GO:0002684 | positive regulation of immune system process |
| GO:0007416 | synapse assembly |
| GO:0070588 | calcium ion transmembrane transport |
| GO:0050776 | regulation of immune response |
| GO:0007218 | neuropeptide signaling pathway |
| GO:0050778 | positive regulation of immune response |
| GO:0016192 | vesicle-mediated transport |
| GO:0016197 | endosomal transport |

| Antenna A | |
|---|---|
| GO:0006508 | proteolysis |
| GO:0006487 | protein N-linked glycosylation |
| GO:0019751 | polyol metabolic process |
| GO:0007112 | male meiosis cytokinesis |
| GO:0006805 | xenobiotic metabolic process |
| GO:0000910 | cytokinesis |
| GO:0010628 | positive regulation of gene expression |
| GO:0033227 | dsRNA transport |
| GO:0010876 | lipid localization |
| GO:0006164 | purine nucleotide biosynthetic process |

| Gut A | |
|---|---|
| GO:0040003 | chitin-based cuticle development |
| GO:0005975 | carbohydrate metabolic process |
| GO:0006508 | proteolysis |
| GO:0006869 | lipid transport |
| GO:0042742 | defense response to bacterium |
| GO:0030001 | metal ion transport |
| GO:0007155 | cell adhesion |
| GO:0009615 | response to virus |
| GO:0008643 | carbohydrate transport |

| Leg N3 | |
|---|---|
| GO:0040003 | chitin-based cuticle development |
| GO:0007591 | molting cycle, chitin-based cuticle |
| GO:0018958 | phenol-containing compound metabolic process |
| GO:0045752 | positive regulation of Toll signaling pathway |
| GO:0016079 | synaptic vesicle exocytosis |
| GO:0070887 | cellular response to chemical stimulus |
| GO:0000422 | autophagy of mitochondrion |
| GO:0008643 | carbohydrate transport |
| GO:0034219 | carbohydrate transmembrane transport |
| GO:0008063 | Toll signaling pathway |

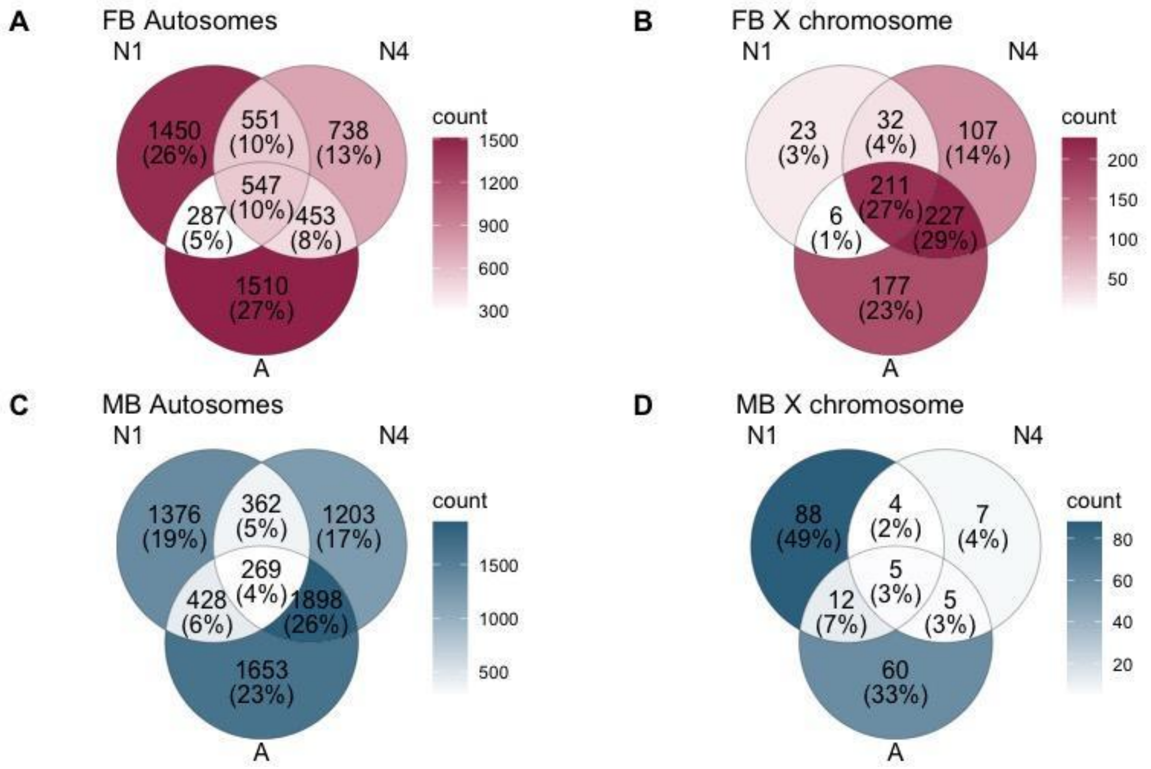


**Supplemental Figure S1.** Venn diagrams showing the overlap of sex-biased genes in the reproductive tract across different developmental stages (A) Female-biased genes on autosomes and on the X chromosome (B) and ale-biased genes on autosomes (C) and on the X chromosome (D)

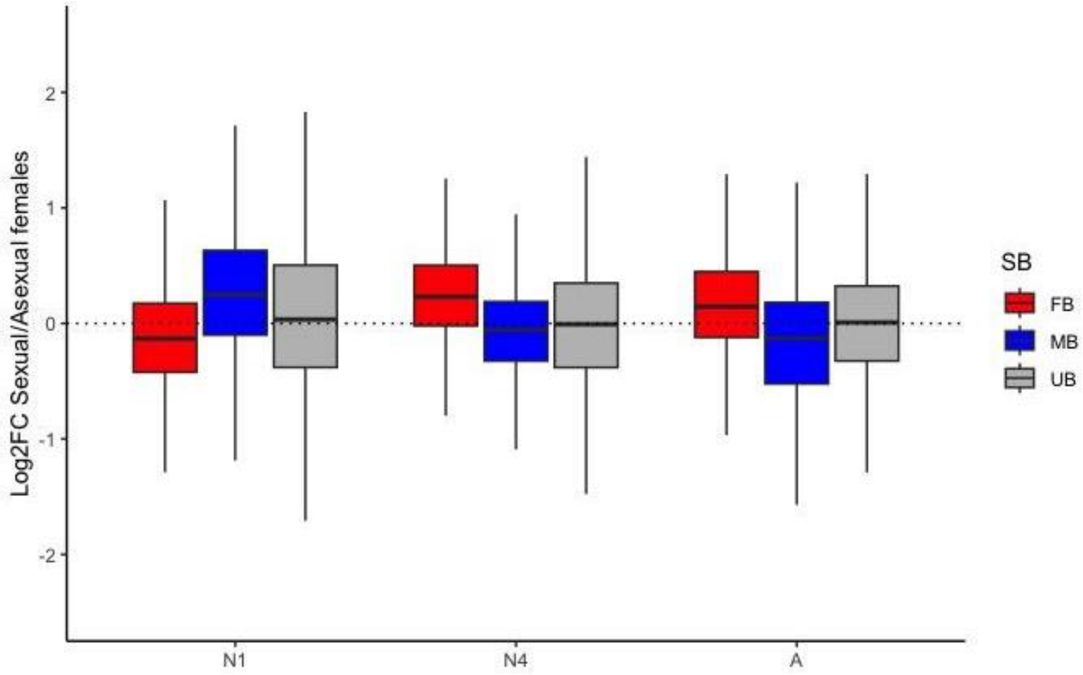


**Supplemental Figure S2.** Shifts in expression levels of sex-biased genes between sexual and parthenogenetic females across development (in reproductive tracts) for genes with low coefficient of variation. Three gene categories are depicted with different colors; female-biased in red, male-biased in blue, and un-biased in grey. Boxplots represent the median, lower and upper quartiles, and whiskers the minimum and maximum values (in the limit of 1.5x interquartile range).

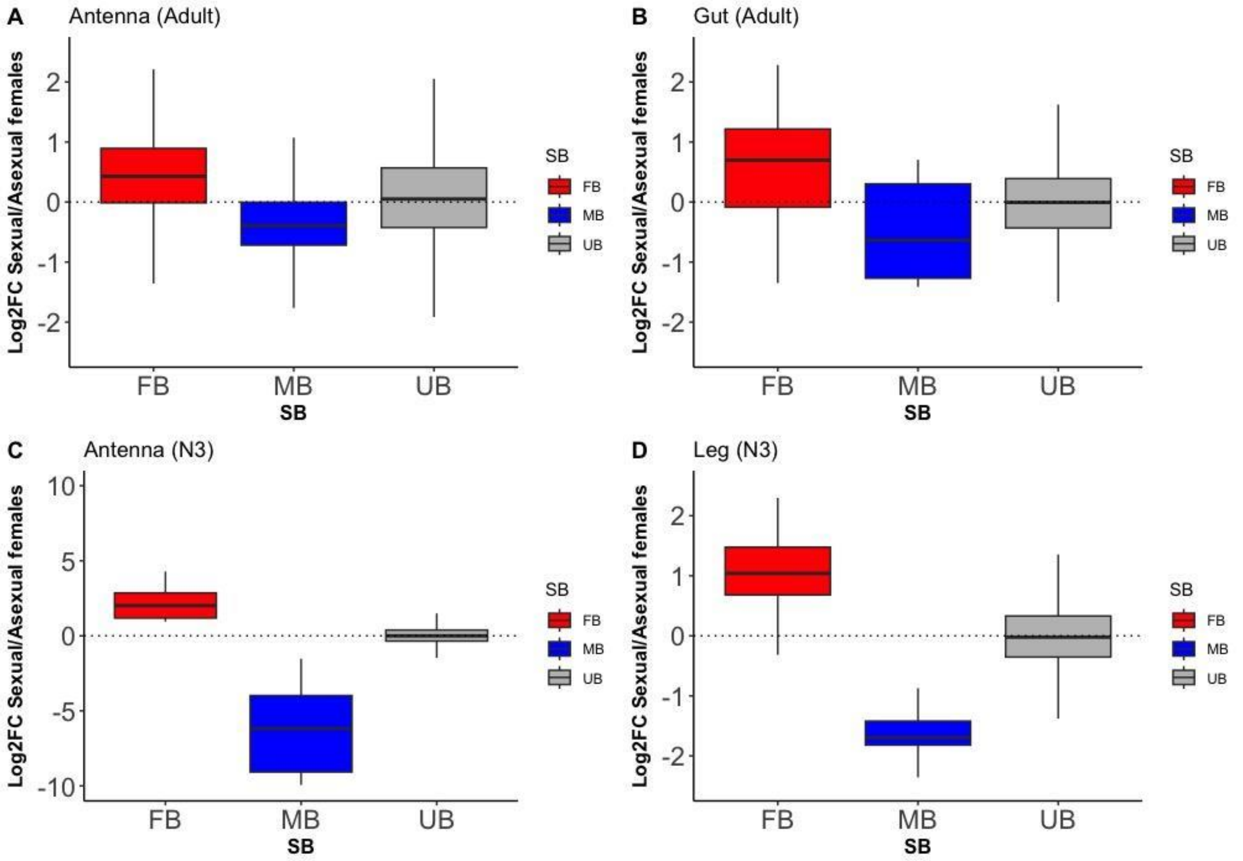


**Supplemental Figure S3.** Shifts in expression levels of sex-biased genes between sexual and asexual females in; A) Antennae- adult stage, B) Gut- adult stage C) Antennae- 3rd nymphal stage and D) Leg- third nymphal stage. Three gene categories are depicted with different colors; female-biased in red, male-biased in blue, and un-biased in grey. Positive values indicate greater expression in parthenogenetic females, negative values indicate greater expression in sexual females. Boxplots represent the median, lower and upper quartiles, and whiskers the minimum and maximum values (in the limit of 1.5x interquartile range). All comparisons between gene categories at each developmental stage yielded statistically significant results, as indicated by adjusted p values from Wilcoxon rank sum tests, with $p < 0.01$.
